# Developing *in vivo* assays for investigation of p75^NTR^ and NRH1 transmembrane domain cleavage events using zebrafish embryos

**DOI:** 10.1101/2020.05.04.076539

**Authors:** Tanya Jayne, Morgan Newman, Michael Lardelli

**Affiliations:** School of Biological Sciences, University of Adelaide, Adelaide, SA, Australia

## Abstract

γ-secretase is an important protease complex responsible for the cleavage of over 100 substrates within their transmembrane domains. γ-secretase acts in Alzheimer’s disease by cleavage of AMYLOID BETA (A4) PRECURSOR PROTEIN to produce aggregation-prone Amyloid beta peptide. Other γ-secretase substrates such as p75^NTR^ are also relevant to Alzheimer’s disease. How γ-secretase cleavage site specificity is determined is still unclear. A previous study using *Xenopus laevis* to investigate the proteolytic processing of p75^NTR^ and its homolog NRH1 found that transmembrane cleavage of NRH1 was not sensitive to the γ-secretase inhibitor DAPT, suggesting that it is not processed by γ-secretase. To investigate this further, we identified zebrafish orthologues of the genes p75^NTR^ and NRH1 and developed *in vivo* assays to assess cleavage of the resultant p75^NTR^ and Nrh1 proteins. Our observations from these assays in zebrafish are consistent with the *Xenopus laevis* study. Inhibition of γ-secretase by DAPT treatment results in accumulation of uncleaved p75^NTR^ substrate, while cleavage of Nrh1 is not affected. This supports that p75^NTR^ is cleaved by γ-secretase while Nrh1 is cleaved by a separate γ-secretase-like activity. We extended our approach by generating a chimeric Nrh1 protein in which the Nrh1 transmembrane domain was replaced by that of p75^NTR^, in an attempt to determine whether it is the p75^NTR^ TMD that confers susceptibility for γ-secretase cleavage. Our results from analysis of this chimeric protein revealed that the p75^NTR^ transmembrane domain alone is insufficient to confer γ-secretase cleavage susceptibility. This is not completely unexpected, as there is evidence to suggest that other factors are crucial for selection/cleavage by the γ-secretase complex. We have established a system in which we can now attempt to dissect the structural basis for γ-secretase cleavage specificity and evolution.

## Introduction

γ-secretase is a multi-subunit membrane-bound aspartyl protease complex responsible for cleavage of over 100 substrates including Amyloid Precursor Protein (APP), Notch and the p75 neurotrophin receptor (p75^NTR^) [1]. γ-secretase has been identified as a member of the intramembrane cleaving protease family (I-CLiP). I-CLiPs cleave type 1 membrane proteins enzymatically via a process termed regulated intramembrane proteolysis (RIP) [2, 3]. The most studied function of γ-secretase is processing of APP. This is due to the seemingly critical role of APP in Alzheimer’s disease (AD) etiology. Although numerous publications have discussed the perceived role of γ-secretase in AD, the specific nature of substrate selection by this protease is not yet clearly defined.

γ-secretase substrates are typically derived from large precursor proteins that undergo a prerequisite removal of their ectodomain/lumenal domain prior to γ-secretase cleavage [3]. Early research suggested that the only other prerequisite was that the substrate must be a type 1 transmembrane protein [4]. Later studies showed that additional factors may guide substrate selection by γ-secretase. It has previously been suggested that dimerisation of substrates and/or the structure of substrate α-helices may regulate γ-secretase activity [5]. Also, γ-secretase substrate recognition and cleavage is much more efficient for ectodomains with fewer than ~50 remaining amino acid residues [6]. Previous studies have attempted to define the amino acid residues of Notch and APP required for γ-secretase cleavage [7]. However, a distinct cleavage recognition site for γ-secretase within the transmembrane domains (TMDs) of its target proteins has not been defined [2].

p75^NTR^, also known as the ‘low-affinity nerve growth factor receptor’ (LNGFR), is one of the many substrates of γ-secretase subject to cleavage within its transmembrane domain [8]. p75^NTR^ has been implicated in neuronal survival, myelination and neurite outgrowth among other pathways during vertebrate nervous system development, through its interactions with neurotrophins and Trk receptors [9]. Also, the Aβ peptide can act as a ligand for p75^NTR^ and is proposed to play a role in cholinergic neuron loss, implicating this protein in AD [10, 11]. Kanning et al (2003) investigated the proteolytic processing of p75^NTR^, along with what they described as the “neurotrophin receptor homologs” (NRH), NRH1 and NRH2 [8]. Database and EST searches have established that the genes coding for these two proteins show greater sequence similarity to p75^NTR^ than to any other homologous genes [8]. Experiments by Kanning *et al* (2003) confirmed processing of p75^NTR^ by α-secretase (ADAM10) and γ-secretase. Western blot analysis of NRH1 and NRH2 indicated that both proteins are cleaved within their transmembrane domains [8]. However, Kanning *et al* (2003) observed that the commonly used inhibitor of γ-secretase activity, DAPT, had no effect on the cleavage of NRH1 or NRH2, suggesting that transmembrane cleavage of these proteins is not by γ-secretase [8].

The possible lack of sensitivity to γ-secretase inhibitors of the p75^NTR^ homologs NRH1 and NRH2 may have interesting applications. p75^NTR^ and its homologs share high sequence similarity in their transmembrane domain, where γ-secretase cleavage occurs [8] and comparison of the transmembrane domains of p75^NTR^ and NRH1 might allow definition of transmembrane domain characteristics critical to permit cleavage by γ-secretase. We have developed an *in vivo* zebrafish assay that can be used to investigate the structural differences in the transmembrane domains of these two proteins that cause their differential sensitivity to γ-secretase. Our results support the observations made by Kanning *et al* (2003), that p75 NTR is cleaved by γ-secretase while Nrh1 is not [8]. We also designed a chimeric construct in which the Nrh1 transmembrane domain is replaced by the TMD of p75^NTR^, to test whether this domain alone can confer γ-secretase cleavage susceptibility to Nrh1. Our results indicate that the p75^NTR^ transmembrane domain alone is insufficient to confer γ-secretase susceptibility to Nrh1.

## Materials and Methods

### Ethics

This work was conducted under the auspices of the Animal Ethics Committee of the University of Adelaide and in accordance with EC Directive 86/609/EEC for animal experiments and the Uniform Requirements for manuscripts submitted to Biomedical journals.

### Gene orthologue identification

Alignments and tree building were conducted using the Geneious software suite, version 5.6.7 (http://www.geneious.com, [12]). Alignments were performed with the following constraints: Cost matrix: Identity, Gap open penalty: 10, Gap extension penalty: 3, Alignment: Global. Bayesian trees were produced using the “Mr Bayes” program with the following constraints: Substitution model: GTR, Outgroup: Lancelet p75^NTR^, and the rest as default.

### Constructs

ssFLAG-p75^NTF^C201-dGFP-v2a-mCherry, ssFLAG-Nrh1C191-dGFP-v2a-mCherry and ssFLAG-A2C-dGFP-v2a-mCherry DNA sequences were produced by Biomatik (complete DNA sequences are provided in text file S1). These DNA sequences in the pBMH vector (provided by Biomaitk) were digested using *Bam*HI and *Cla*I in independent reactions and ligated into pT2AL200R150G between the *Bam*HI and *Cla*I sites.

### Replacement of mCherry coding sequence by GFP coding sequence

GFP was amplified by polymerase chain reaction (PCR) using the following primers; Forward: 5’-GCTCTAGAATGGTGAGCAAGGGAGAGGA-3’ and Reverse: 5’CCATCGATCTACTTGTACAGCTCGTCCATTCC-3’. The thermal cycling parameters were as follows: 98°C for 30 s, 15 cycles of 98°C for 10 s, 61°C for 30 s and 72°C for 30 s, followed by 72°C for 10 mins. mCherry was excised from pT2AL200R150GssFLAG-p75^NTF^C201-dGFP-v2a-mCherry, pT2AL200R150G ssFLAG-Nrh1C191-dGFP-v2a-mCherry and pT2ALssFLAG-A2C-dGFP-v2a-mCherry by restriction digest with *Xba*1 and *Cla*1. The amplified GFP was then cloned between the *Xba*1 and *Cla*1 sites of pT2AL200R150GssFLAG-p75^NTF^C201-dGFP-v2a-mCherry, pT2AL200R150G ssFLAG-Nrh1C191-dGFP-v2a-mCherry and pT2ALssFLAG-A2C-dGFP-v2a-mCherry.

### DNA microinjection of zebrafish embryos and treatment with DAPT

Tol2 transposase plasmid (pCS-TP) was linearised using Not1 (NEB) and mRNA was transcribed *in vitro* using the mMESSAGE mMACHINE SP6 Kit (Ambion Inc.). Fertilised embryos were injected with a solution containing 100ng/µl plasmid DNA and approximately 50ng/µl Tol2 transposase mRNA at the one cell stage. ~50 embryos injected with the injection solution above were placed into 35mm × 10mm petri dishes with 2ml E3 medium (15mM NaCl, 0.5mM KCl, 1mM MgSO_4_, 0.15mM KH_2_PO_4_, 0.05mM Na_2_HPO_4_, 1mM CaCl_2_, 0.7mM NaHCO_3_). At 4 hours post fertilisation (hpf) embryos were treated with 100µM DAPT (In solution™ γ-secretase inhibitor IX, Calbiochem, San Diego, CA, USA) in 1% DMSO in E3 medium. Embryos were maintained at 28°C in a humid incubator. At 24 hpf embryos were visualised under UV light for GFP expression. Embryos expressing GFP were selected for protein extraction.

### Treatment of zebrafish embryos with Z-DEVD-FMK

Injected embryos were placed into 35mm × 10mm petri dishes with 2mL E3 medium (15mM NaCl, 0.5mM KCl, 1mM MgSO4, 0.15mM KH2PO4, 0.05mM Na2HPO4, 1mM CaCl2, 0.7mM NaHCO3). At 6 hpf embryos were treated with 100µM Z-DEVD-FMK (ApexBio Technology LLC, Houston, TX, USA) in E3 medium. Embryos were maintained in standard temperature conditions in a humid incubator and checked for survival at 24 hpf.

### Western Immunoblot analyses

Dechorioned and de-yolked embryos were lysed by placement in sample buffer (2% sodium dodecyl sulfate (SDS), 5% β-mercaptoethanol, 25% v/v glycerol, 0.0625 M Tris-HCl (pH 6.8), and bromophenol blue) followed immediately by heating to 95°C for 10 min, before storage at −80°C prior to protein separation on 4-12% SDS polyacrylamide gels. Proteins were transferred to nitrocellulose membrane in buffer (25mM Tris, 192mM glycine, 0.1% sodium lauryl sulfate, 20% methanol in MilliQ H_2_O) at 10V for 1hr. When immunoblotting, all membranes were blocked with 5% Western Blocking Reagent (Roche, Indianapolis, IN, USA) in TBST, incubated with primary antibodies in TBST containing 0.5% Western Blocking Reagent (Roche, Indianapolis, IN, USA), washed in TBST, and incubated in secondary antibody. Following secondary antibody incubations, all membranes were washed three times for 10 minutes in TBST and visualised with luminol reagents (Amresco, Ohio, USA <u>or</u> Thermo Scientific, Rockford, USA) by the ChemiDoc™ MP imaging system (Bio-Rad, Hercules, CA, USA). The p75^NTF^C201-dGFPx2, Nrh1C191-dGFPx2 and A2C-dGFPx2 protein bands were visualised at ~61kDa, 57kDa and 57kDa respectively. Using Image Lab software (Bio-Rad), densitometry analyses were performed on the protein bands in each lane of the western immunoblot for each sample and free GFP internal reference. An average value was obtained for the free GFP for each membrane. Each sample value was then normalised to the average free GFP value.

GFP immunoblots were incubated in a 1/5,000 dilution of anti-GFP antibodies (Rockland Immunochemicals Inc., Gilbertsville, PA, USA) and a 1/10,000 dilution of donkey anti-Goat IgG (Rockland Immunochemicals Inc., Gilbertsville, PA, USA).

mCherry immunoblots were incubated in a dilution of 1/2,000 of anti-mCherry antibody (Abcam, Cambridge, UK) and a 1/5,000 dilution of anti-Mouse IgG secondary antibodies (Rockland Immunochemicals Inc., Gilbertsville, PA, USA).

### Statistical analyses

Unpaired, two-tailed, t-tests with Welch’s correction (assuming Gaussian distribution), were performed on normalised values using GraphPad Prism version 8.0.0 for Windows, GraphPad Software, San Diego, California USA, www.graphpad.com. P values of < 0.05 were considered to be significant. We did not remove any outliers from the analysis.

## Results and Discussion

### Identification of p75^NTR^ and NRH1gene orthologues in zebrafish

Of the two *NRH* genes, *NRH1* has coding sequences more similar in length to *p75^NTR^* and is the only know *NRH* gene with a putative orthologue in zebrafish. To investigate p75^NTR^ and NRH1 cleavage *in vivo* using zebrafish, we first validated the orthology of these genes. A tblastn search performed against the zebrafish genome using the entire putative protein sequence of human *p75^NTR^*, constrained to “RefSeq_RNA”, returned candidate orthologues of the human *p75^NTR^* gene on zebrafish chromosomes 3, 12 and 16. At the time of the original analysis, chromosome 12 appeared to hold two almost identical copies of the gene at different loci (supplementary file S1, Figure S1) which we suspected was due to a recent duplication event. The position of the duplicate appears to have been revised to chromosome 3 in the latest genome build (GRCz11) (Figure S2). The top tblastn hit, *nerve growth factor receptor b* (*ngfrb*), on chromosome 12, has the greatest query cover (percentage of the sequence aligned to a sequence in GenBank) to human *p75^NTR^* (93%) (Figure S2), so we tentatively named this “zebrafish p75”. A tblastn search performed against the zebrafish genome using *NRH1* from *Xenopus laevis* (GenBank accession AF131890.1) returned the computer predicted sequence for *neurotrophin receptor associated death domain* (*nradd*) on chromosome 16 (also returned as a best hit in the human *p75^NTR^* tblastn search described above) with 100% query coverage to Xenopus *NRH1*. The only other strong zebrafish *Nrh1* candidate returned was *ngfrb*, which we had already established most likely represents *p75^NTR^* in zebrafish. Therefore, we predict that *nradd* is most likely a *Nrh1* orthologous gene in zebrafish.

NRH1 belongs to a subfamily of vertebrate p75^NTR^-related proteins which also contains NRH2. NRH2 exists only in mammals while NRH1 exists only in amphibians, fish and birds [8]. The return of the predicted sequence *nradd* when searching for *NRH1* within the zebrafish genome is consistent with previous knowledge that *NRH2* is also known as *NRADD* in mouse and rat (mouse NCBI Gene ID: 67169, rat NCBI Gene ID: 246143). Therefore, we eliminated *nradd* as a zebrafish p75^NTR^ candidate gene and propose that it is the *Nrh1* orthologous gene in zebrafish.

To confirm our identification of zebrafish *p75^NTR^* and *Nrh1* orthologues we next conducted phylogenetic analyses using the Geneious software suite [12]. Zebrafish p75^NTR^ and Nrh1 candidate amino acid sequences along with p75^NTR^ amino acid sequences from *Xenopus laevis*, *Gallus gallus* (chicken), mouse and human and NRH1 amino acid sequences from *Xenopus tropicalis*, *Xenopus laevis* and chicken were aligned using the Geneious alignment tool, and trees were built using both Bayesian and Maximum likelihood methods. We included the amino acid sequences of NRH2 from mouse and human in these analyses as these are the mammalian equivalents of NRH1 [8]. Accession numbers for all sequences used can be found in Table S1. *Branchiostoma floridae* (lancelet) was used as an out-group as this was the most distant relative to zebrafish that returned a result when conducting tblastn searches using human *p75^NTR^* and *X. laevis NRH1*. Interestingly, tblastn searches of the lancelet genome using both *p75^NTR^* and *NRH1* returned the same gene in lancelet (Table S1). As the chicken genome contains both *p75^NTR^* and *NRH1*-like sequences, tblastn searches of the lancelet genome using both full-length chicken sequences were performed to confirm the preliminary findings. These searches returned results identical to those using human *p75^NTR^* and *X. laevis NRH1*. This supports that there is only a single *p75^NTR^*- and *NRH1*-like gene in this basal chordate and that *p75^NTR^* and *NRH1* arose from a gene duplication event early in vertebrate evolution.

A dendrogram modelling the phylogenetic relationships of p75^NTR^ and its homologs demonstrated p75^NTR^ proteins clustering together and NRH1 proteins clustering together in separate clades (Fig 1). This supports that the sequence *nradd* on chromosome 16 of zebrafish is indeed the orthologue of *Nrh1*.

**Fig 1.**
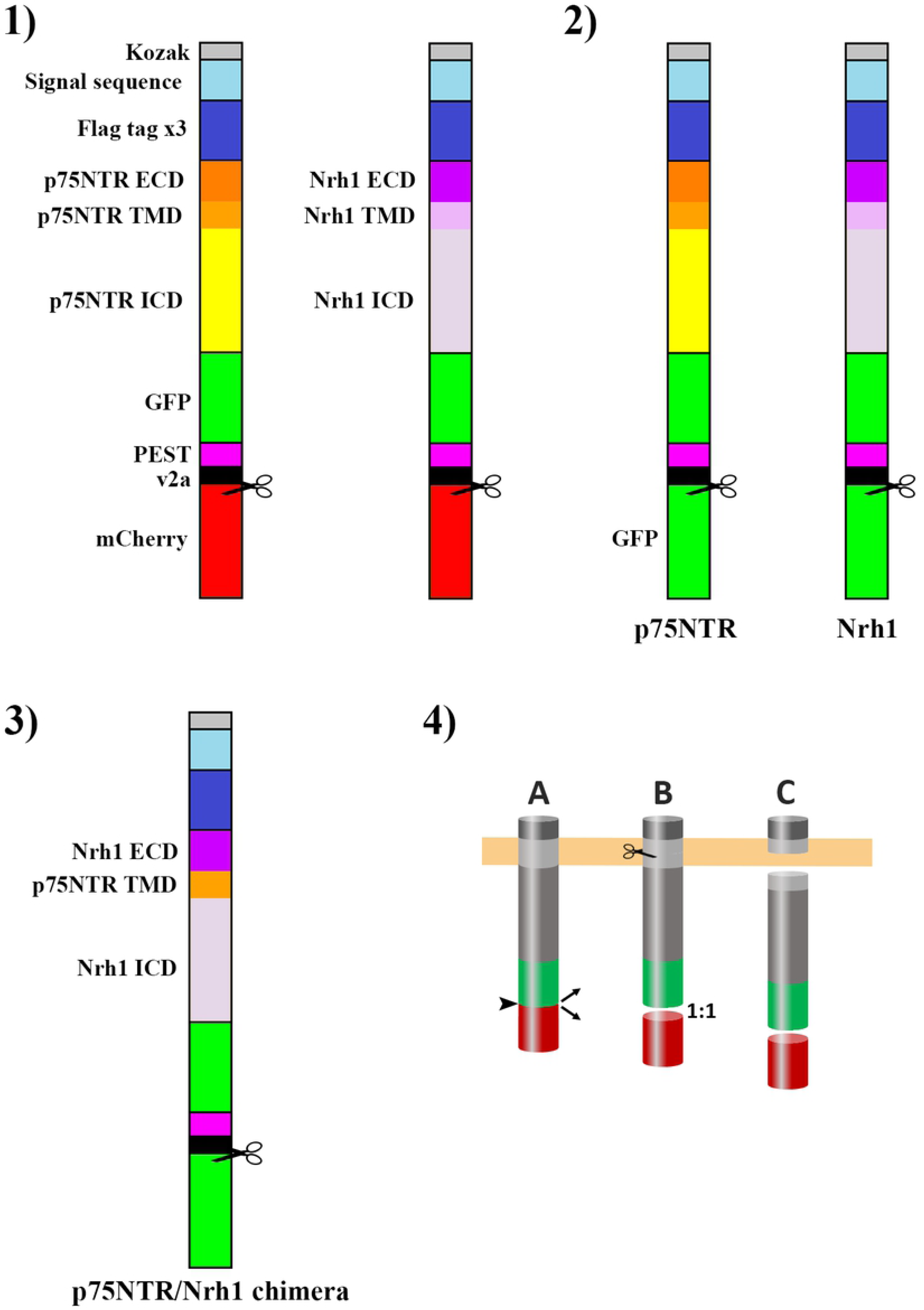
Phylogenetic tree of p75^NTR^ and NRH1 proteins. This tree was generated using the Geneious software suite to perform Bayesian analysis (MrBayes). Numbers represent the aLRT branch-support values.

It was difficult to discern which of the remaining three possible zebrafish p75^NTR^ protein sequences was most likely to represent the true p75^NTR^ in zebrafish from the phylogenetic analyses, although the sequence that we designated zebrafish p75-like-chr3 appeared to be marginally more similar to human p75^NTR^ (Fig 1). The sequence we designated zebrafish p75, known as *ngfrb* in the NCBI database, is the only gene with a coding sequence not derived by computer-prediction from genomic DNA sequence. It was thereby judged to be the most likely to be accurate and was selected for construction of an assay.

### Design of p75^NTR^ and Nrh1 γ-secretase cleavage assay constructs

In order to be recognised as a cleavage substrate by γ-secretase, proteins must first undergo truncation of the extracellular domain [6]. This preliminary shedding of large extracellular domains by α- or β-secretase may be rate-limiting, creating problems for an *in vivo* assay as the rate of cleavage may be dependent on these events rather than γ-secretase itself. To overcome this in our assay, we proposed to produce artificially truncated forms of both proteins similar to a strategy we had adopted previously for zebrafish APP in another γ-secretase assay [13].

To truncate p75^NTR^ and Nrh1 to mimic approximately α-secretase-cleaved forms of these proteins, we first needed to identify the transmembrane domains of both p75^NTR^ and Nrh1. As the transmembrane domains of zebrafish p75^NTR^ and Nrh1 genes had not yet been defined, we employed online prediction programs “TMHMM server, v. 2.0” (http://www.cbs.dtu.dk/services/TMHMM/) [14], “TMpred Server” (https://embnet.vital-it.ch/software/TMPRED_form.html) and “DAS” (https://tmdas.bioinfo.se/DAS/index.html, [15]) to determine the location of the transmembrane region within each protein (supplementary data File S3, Figure S3). The success of this approach was confirmed by comparing our predicted transmembrane domains to the defined transmembrane domain of human p75^NTR^ [5]. It has previously been established that γ-secretase cleavage of p75^NTR^ is dependent on shedding of the N-terminal extracellular domain by the α-secretase, A DISINTEGRIN AND METALLOPROTEINASE DOMAIN 17 (ADAM17). A previous study of p75^NTR^ suggested that this cleavage occurs five amino acids before the N-terminal of the transmembrane domain [5]. Another study found that a large deletion of 15 amino acid residues within the juxtamembrane domain (before the N-terminal of the transmembrane) of p75^NTR^ led to substantial decrease in shedding, while a third study found that a stub of 15 amino acids N-terminal to the transmembrane domain was sufficient for γ-secretase cleavage [16, 17]. We elected to remove most of the extracellular domain, leaving only 15 amino acids immediately adjacent to the N-terminal end of the transmembrane domain. The C-terminal intracellular domain was left at its original length as it has not been found to affect cleavage [16]. When attempting to visualise cleaved and uncleaved protein fragments on a western blot this small 15 residue N-terminal stub may have posed a problem. After γ-secretase cleavage we would be left with a very short N-terminal fragment and a much larger C-terminal fragment, which would presumably be difficult to resolve on the same western immunoblot.

To overcome this, 3 FLAG tags [18] were fused in tandem to the N-terminals of both p75^NTR^ and Nrh1. The efficiency of processing by γ-secretase is not reduced unless the number of amino acids in the extracellular domain exceeds 50, so processing and cleavage should not be affected by the FLAG tags [6]. Single, silent, point mutations were introduced into both the second and third tandem FLAG tag repeats to inhibit recombination in bacteria during cloning [19]. The fourth codon in the second repeat was altered from GAT to GAC, and the sixth codon in the third from GAC to GAT respectively. The highly active HMM+38 secretory signal sequence was added N-terminal to the FLAG repeats to ensure insertion of the p75^NTR^ and Nrh1 proteins into lipid bilayers [20]. This signal sequence is cleaved off upon reaching the target site and is not involved in the metabolism of the final translated protein, hence does not alter the mature protein structure [20]. Destabilised green fluorescent protein (dGFP) was included at the C-terminus of p75^NTR^ and Nrh1 to allow for visualisation of expression *in vivo* and via western immunoblot analysis. As these constructs contain a truncated, C-terminal fragment of the original p75^NTR^ and Nrh1 proteins we describe them as ssFLAG-p75^NTR^C201-dGFP and ssFLAG-Nrh1C191-dGFP respectively.

Injection of transposon-based vector DNA into fertilised zebrafish eggs can give variable results in terms of the amount of DNA delivered and the subsequent degree of transposition and transgene expression. Therefore, when examining the stability of a protein expressed from an injected transgene, it can be useful to have an internal reference standard against which the quantity of the protein can be compared. In a previous study assaying γ-secretase activity by monitoring cleavage of zebrafish Appa fused to dGFP (also expressed from the Tol2 transgene vector, pT2AL200R150G), we co-injected a similar Tol2 vector expressing free GFP, as an internal reference standard [13]. However, results from strategy may be prone to variability as the free GFP vector might not transpose into the genome at a constant rate relative to the Appa construct.

The viral 2A (v2A) peptide ribosomal-skip mechanism allows for expression of two different proteins independently of one another. The skip mechanism occurs within the v2A sequence when a peptide bond fails to form between the penultimate (glycine) and final (proline) residue. Translation continues despite this failure, and tandem protein products are produced in a stoichiometric manner [21]. In order to express truncated p75^NTR^ or Nrh1 simultaneously with an internal reference standard from the same expression vector, we included a v2a sequence at the C-terminal of ssFLAG-p75^NTR^C201-dGFP and ssFLAG-Nrh1C201-dGFP followed by coding sequence for the red fluorescence protein, mCherry. We describe these Tol2-based expression constructs as pT2ALssFLAG-p75^NTF^C201-dGFP-v2a-mCherry and pT2ALssFLAG-Nrh1C191-dGFP-v2a-mCherry (Fig 2, 1). For simplicity, we will henceforth refer to them as p75^NTF^C201-dGFP and Nrh1C191-dGFP respectively. This design enables stoichiometric production of p75^NTF^C201-dGFP or Nrh1C191-dGFP simultaneously with mCherry, allowing for normalisation of protein expression between successive batches of injected embryos.

**Fig 2.**
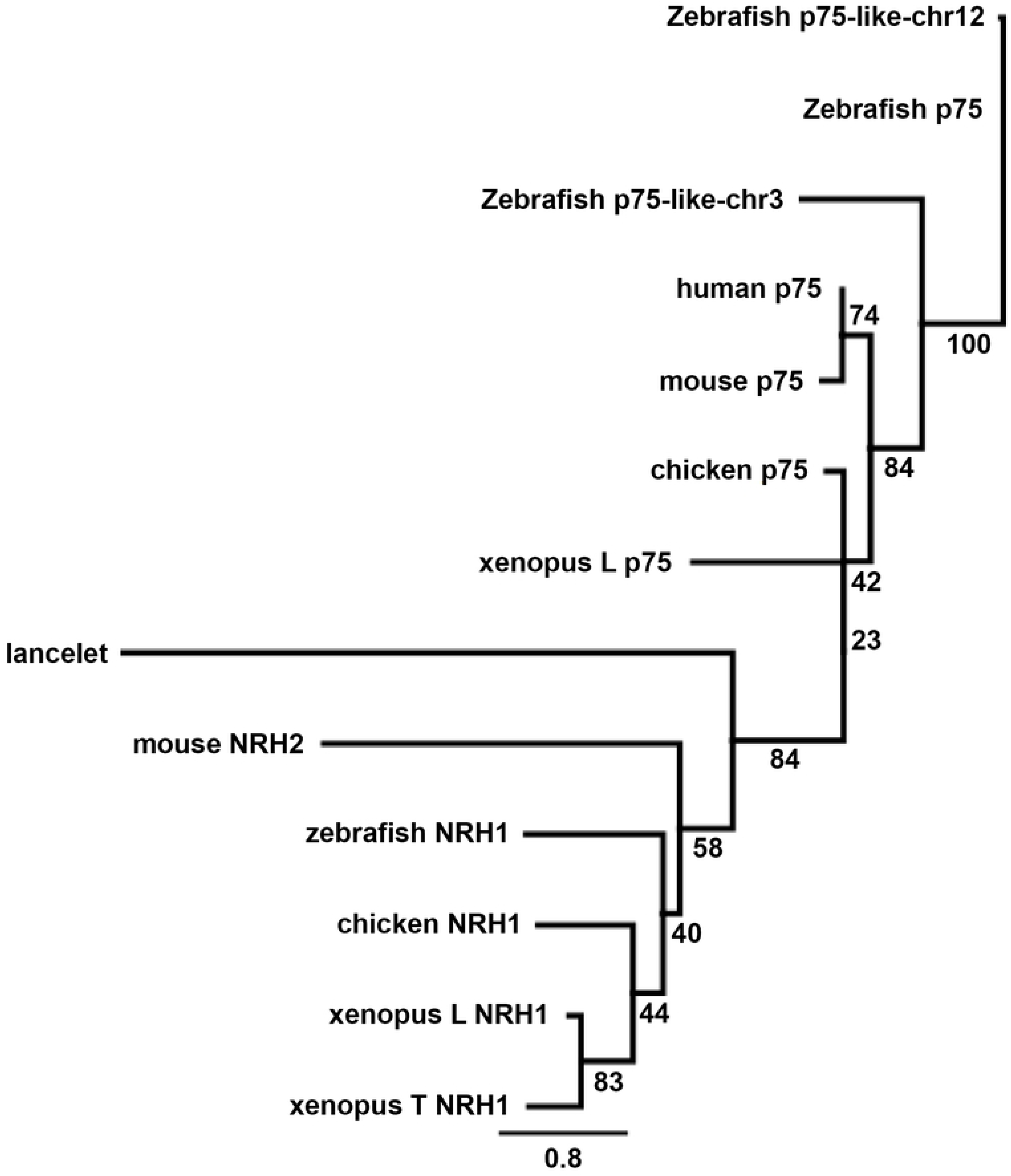
Overview of all construct designs and assay process. 1) Overview of pT2ALssFLAG-p75^NTF^C201-dGFP-v2a-mCherry and pT2ALssFLAG-Nrh1C191-dGFP-v2a-mCherry constructs. 2) pT2ALssFLAG-p75^NTF^C201-dGFP-v2a-GFP and pT2ALssFLAG-Nrh1C191-dGFP-v2a-GFP. 3) Overview of pT2ALssFLAG-A2C-dGFP-v2a-GFP. 4) Anticipated assay process. A) the position and mechanism of the ribosomal skip is indicated by arrows, B) TMD cleavage of constructs is represented by scissors. In all figures ECD = extracellular domain, TMD = transmembrane domain and ICD = intracellular domain.

To investigate γ-secretase cleavage of both p75^NTF^C201-dGFP and NRH1C191-dGFP, each expression vector was co-injected with transposase mRNA into single-cell stage zebrafish embryos. At 24 hours post fertilisation (hpf) embryos displaying GFP fluorescence under UV light were selected, their yolks removed, and protein extracted by lysis in SDS buffer. Protein lysates were separated by SDS polyacrylamide gel electrophoresis and then subjected to western immunoblotting to detect GFP. Uncleaved p75^NTF^C201-dGFP and Nrh1C191-dGFP proteins were visible as bands of ~61kDa and ~57kDa respectively and could be assessed by densitometry, while their intra-membrane domain cleavage products could not be observed (presumably due to their instability). Stripping and probing of the blot to detect mCherry followed by densitometry allowed normalisation of the GFP signals to facilitate comparison of p75^NTF^C201-dGFP and Nrh1C191-dGFP stability between samples.

### Investigating differential cleavage properties of zebrafish p75^NTR^ and Nrh1

Previous experiments performed by Kanning *et al* (2003) indicated that both the p75^NTR^ and NRH1 proteins are cleaved within their transmembrane domains. However, treatment with the known γ-secretase inhibitor DAPT had no effect on NRH1 cleavage, suggesting that NRH1 is not processed by γ-secretase [8]. To test whether p75^NTF^C201-dGFP and Nrh1C191-dGFP cleavage displayed differential sensitivity to inhibition of γ-secretase activity *in vivo*, batches of injected embryos were divided into two groups. Half of a batch injected with either p75^NTF^C201-dGFP or Nrh1C191-dGFP was treated with 100μM of the γ-secretase inhibitor DAPT from 4 hpf until 24 hpf, while the other half was left untreated. DAPT is a potent γ-secretase cleavage inhibitor, hence we expected that treatment with DAPT would result in an accumulation of uncleaved substrates of γ-secretase [17]. This was observed for p75^NTF^C201-dGFP-injected embryos treated with DAPT (Fig 3, A), suggesting that zebrafish p75^NTR^ is, indeed, processed by γ-secretase. However, when western immunoblot densitometry data was normalised to mCherry across three replicates, the p-value of this observed increase (p = 0.1697) did not support the likelihood that uncleaved zebrafish p75^NTR^ is consistently accumulated when treated with DAPT (Fig 3, B). This accumulation of substrate was not observed when Nrh1C191-dGFP was subjected to DAPT treatment (Fig 3, A), suggesting that zebrafish Nrh1 is not sensitive to the γ-secretase inhibitor DAPT. This result supports the observation made by Kanning *et al* (2003) that NRH1 is not a substrate of γ-secretase [8].

**Fig 3.**
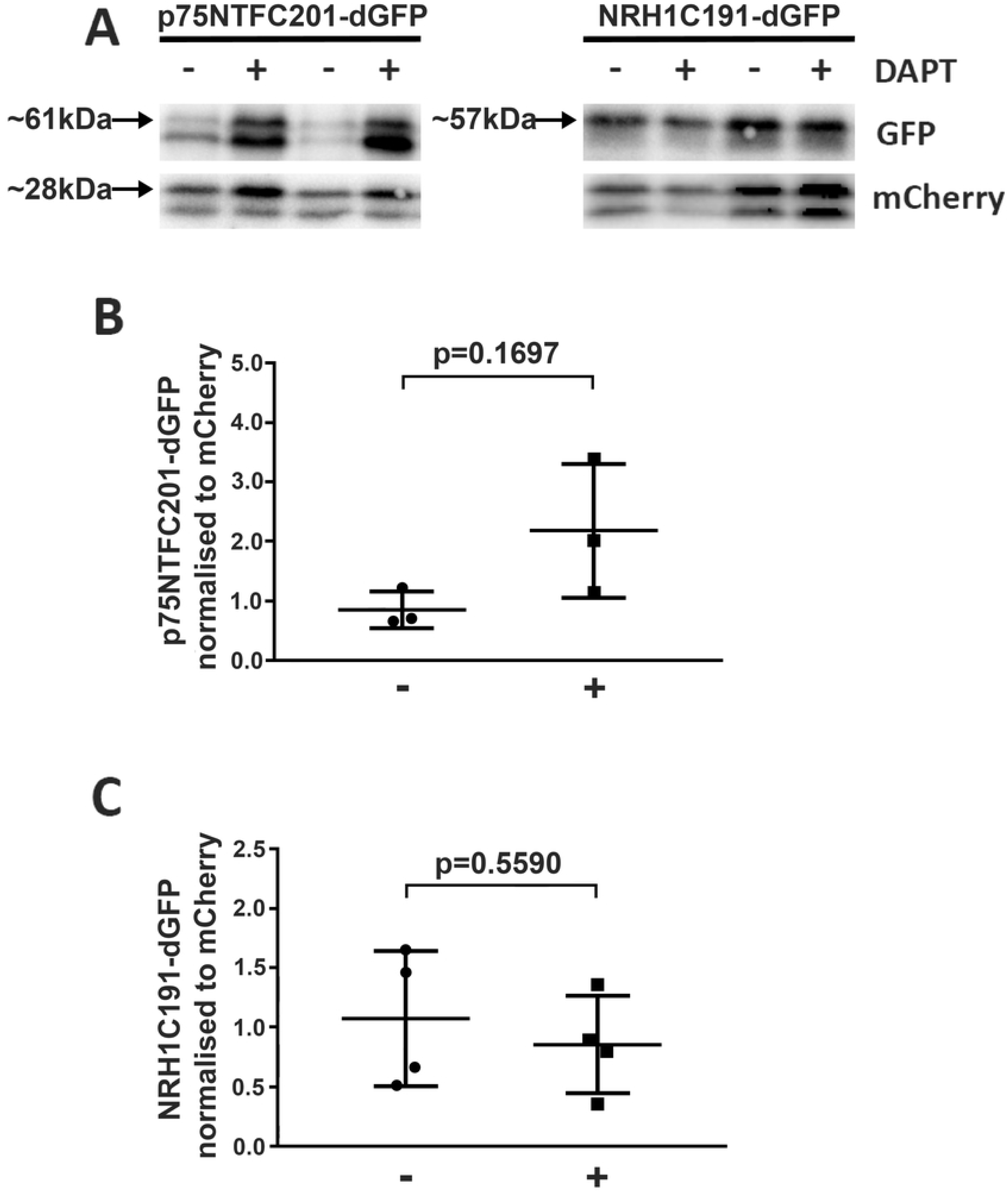
Western immunoblot analysis of p75^NTF^C201-dGFP and Nrh1C191-dGFP. A) Western immunoblots from p75^NTF^C201-dGFP and Nrh1C191-dGFP injected embryos at 24 hpf with and without DAPT treatment. + indicates embryos were treated with DAPT. The additional band observed below the bands indicated by arrows are most likely degradation products. B) Ratios of p75^NTF^C201-dGFP/free mCherry in p75^NTF^C201-dGFP injected embryos at 24 hpf, with (+) (n = 3) or without (-) (n = 3) DAPT treatment. C) Ratios of Nrh1C191-dGFP /free mCherry in Nrh1C191-dGFP injected embryos at 24 hpf, with (+) (n = 4) or without (-) (n = 4) DAPT treatment.

As was previously observed in a similar γ-secretase assay, both p75^NTF^C201-dGFP and Nrh1C191-dGFP appeared to induce developmental abnormalities in embryos [13]. When injected, embryos displayed increased mortality (~46% vs. ~5-10% in uninjected controls) and a range of developmental abnormalities, which were increased by DAPT treatment (~57% mortality). This toxicity may be due to excessive generation of p75^NTF^ and Nrh1 intracellular domains (ICDs), both of which carry death domains [5, 8]. This increase in ICD carrying death domains may result in premature embryo death. p75^NTF^ is also known to form a heterodimer with the protein SORTILIN which then interacts with proNGF or proBDNF proteins leading to apoptosis [22, 23]. Treatment of embryos with DAPT may inadvertently facilitate this interaction by causing accumulation of uncleaved p75^NTR^. Another factor to consider is the nature of the vectors used to express the assay constructs. Tol2 is a transposase vector that inserts randomly into the genome. If the Tol2-based constructs were to disrupt essential/housekeeping genes this would also affect the survival of embryos [24].

Zebrafish caspase 3 plays an important role in apoptosis signalling [25]. To overcome the toxicity of assay construct expression, we treated zebrafish embryos with 100μM of the caspase 3 inhibitor Z-DEVD-FMK (as was shown in a PhD dissertation [26]). However, treatment with this inhibitor did not reduce the degree of lethality observed. Therefore, to overcome the problem of the lethality, an increased number of embryos were injected in each batch with only the most phenotypically normal embryos being selected for western blot analysis (after confirmation that they were expressing GFP observable by its fluorescence).

### Replacing mCherry with GFP reduces variability in the western immunoblot analyses

An apparent trend of accumulation of p75^NTF^C201-dGFP due to γ-secretase inhibition was observed by western immunoblotting. However, statistical analysis of the densitometry measurements did not indicate significance due to the considerable variability between samples (Fig 3, B and C). A contributor to this variability may have been the necessity to strip and re-probe the western blot with the anti-mCherry antibody. To overcome this, the red fluorescence gene mCherry was excised from the constructs and replaced with a second GFP gene downstream of the C-terminal of v2a, producing vectors pT2ALssFLAG-p75^NTF^C201-dGFP-v2a-GFP and pT2ALssFLAG-Nrh1C191-dGFP-v2a-GFP (Fig 2, 2). For simplicity, we will henceforth refer to these as p75^NTF^C201-dGFPx2 and Nrh1C191-dGFPx2 respectively. This minor adjustment in construct design allows for the internal expression standard to be visualised using the same anti-GFP antibody as detects the p75^NTF^C201-dGFP and Nrh1C191-dGFP fusions.

To evaluate the effectiveness of the modified assay constructs we performed injections on numerous batches of embryos and then ran protein samples on multiple western blots. Analysis using the new assay constructs consistently displayed an increase in accumulation of the p75^NTF^C201-dGFPx2 substrate when treated with the γ-secretase inhibitor DAPT. This result was confirmed statistically by combining band intensity data from across all western blots and performing a two-tailed t-test assuming unequal variances, resulting in a p value of 0.0047 (Fig 4, A).

**Fig 4.**
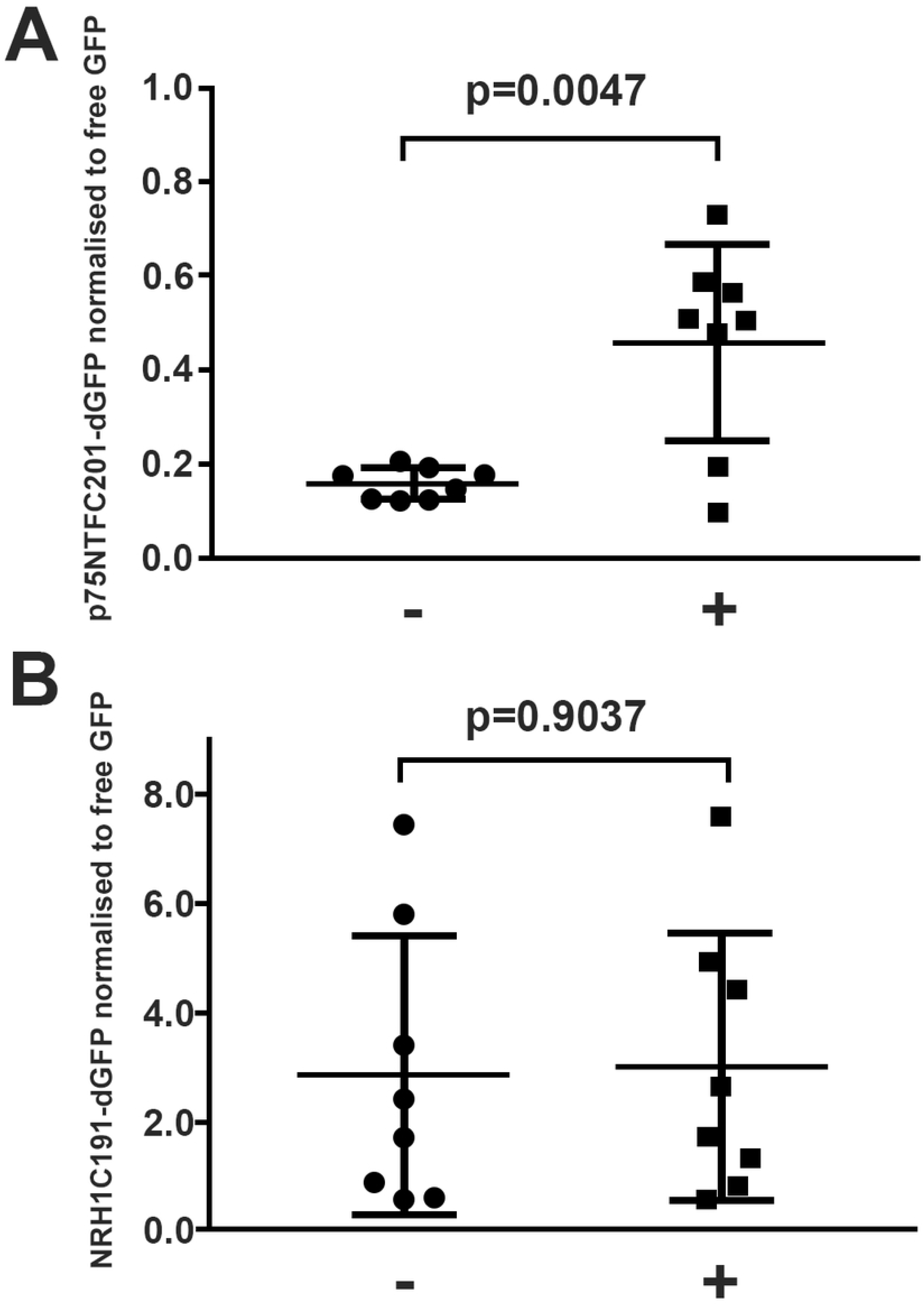
Western immunoblot analysis of p75^NTF^C201-dGFPx2 and Nrh1C191-dGFPx2. Ratios from western immunoblots of p75^NTF^C201-dGFPx2/free GFP in p75^NTF^C201-dGFPx2 injected embryos at 24 hpf, with (+) (n = 8) or without (-) (n = 8) DAPT treatment. C) Ratios of Nrh1C191-dGFPx2 /free GFP in Nrh1C191-dGFPx2 injected embryos at 24 hpf, with (+) (n = 8) or without (-) (n = 8) DAPT treatment. p value was calculated using an unpaired, two-tailed t-test.

Conversely, there was no observed increase in Nrh1C191-dGFP substrate accumulation in response to DAPT treatment. Statistical analysis found no significant difference between the treated and untreated samples (p =0.9037) (Fig 4, B). Although the p75^NTR^ western immunoblot data was similar across numerous blots there was a high degree in variability in the normalised values across Nrh1 immunoblots. This variability was unexpected and seems to be due to variation in the amount of free GFP on each blot. It is possible that GFP stability may play a role in this. Regardless of this we are confident that Nrh1 is not responsive to DAPT as, when looking at each blot individually, there was consistently no accumulation in response to this inhibitor among replicates.

### γ-secretase cleavage is not conferred to Nrh1 by the p75^NTF^ transmembrane domain

The sequence similarity of p75^NTR^ and its homolog Nrh1 imply that these two genes share a relatively recent evolutionary origin through duplication of an ancestral p75^NTR^/Nrh1-like gene. Therefore, if p75^NTR^ is a substrate of γ-secretase, then Nrh1 would most probably also be cleaved by it. The observed lack of γ-secretase-dependent cleavage of zebrafish Nrh1 in our assay is consistent with the results of Kanning *et al* (2003) [8]. The existence of this pair of closely related genes/proteins, one of which is cleaved by γ-secretase and one of which is not, presents us with a unique opportunity to dissect the structural basis of γ-secretase cleavage substrate specificity.

To begin dissection of γ-secretase cleavage substrate specificity using our assay, we designed a chimaeric construct in which Nrh1’s transmembrane domain was replaced with the transmembrane domain from p75^NTR^. The new construct, termed pT2ALssFLAG-A2C-dGFP-v2a-GFP (Fig 2, 3), simplified to A2C-dGFPx2, was injected into one cell stage embryos which were subsequently treated as previously with or without DAPT. Protein samples were then collected at 24 hpf for analysis by western immunoblot. This did not reveal an accumulation of substrate when γ-secretase was inhibited (Fig 5). This suggests that the p75^NTR^ transmembrane domain alone is not sufficient to confer γ-secretase susceptibility and that structures outside of this domain are also required. It is possible that altering the protein by swapping entire domains may disrupt its ability to form homo- or heterodimers, which have previously been found to be important for γ-secretase cleavage [5]. A detailed further analysis replacing Nrh1 amino acid residues with those not shared by p75^NTR^ in this chimaeric construct should allow definition of the structures critical to conferring γ-secretase susceptibility.

**Fig 5.**
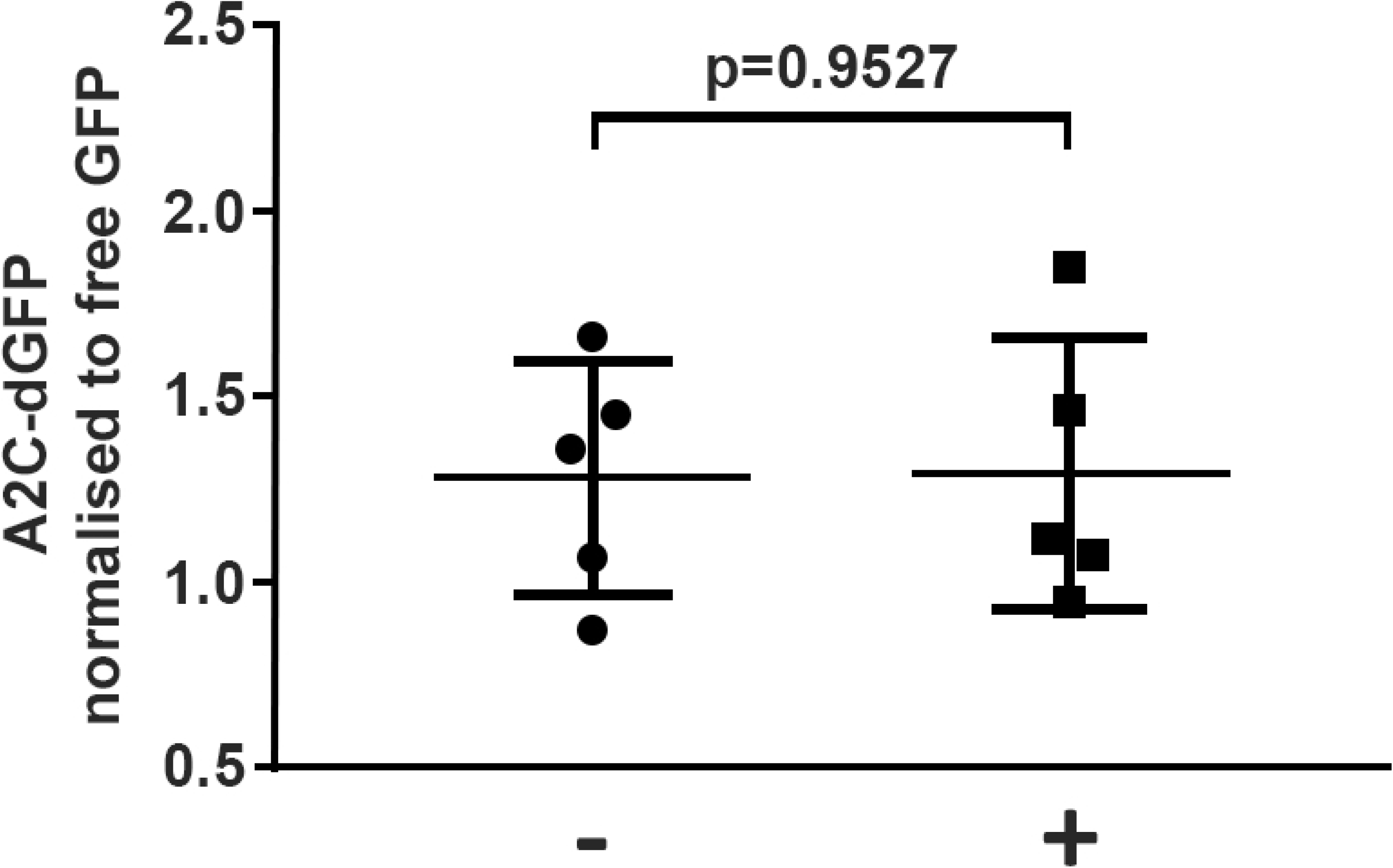
Western immunoblot analysis of A2C-dGFPx2. Ratios from western immunoblots of A2C-dGFPx2 /free GFP in A2C-dGFPx2 injected embryos at 24 hpf, with (+) (n = 5) or without (-) (n = 5) DAPT treatment.

### Further analysis of p75^NTR^ and Nrh1 transmembrane domains

Recent studies have shown that the α-helices of γ-secretase substrates Notch and APP unwind when they interact with the active site of PRESENILIN (the catalytic core of γ-secretase) [27–29]. A study investigating the conformation of the rhomboid substrate Gurken during cleavage by the archaeal homologue of PRESENILIN, MCMJR1, found that Gurken underwent a conformational change into a β-strand when interacting with MCMJR1 [30]. Proline residues have previously been found to disrupt transmembrane helices. Therefore, Brown *et al* (2018) altered the Pro252 TMD residue of Gurken and found that it was no longer a substrate of MCMJR1. They suggested that the proline residue, through its perturbation of the α-helical conformation, effectively allows for this helix to unwind into the β-strand conformation that is preferred by MCMJR1 [30]. They also observed that so called “noncleavable” variants (i.e. those that cannot enter a β-strand conformation) could still bind to MCMJR1 with equal affinity to the cleavable substrates.

We aligned the TMDs of zebrafish p75^NTR^ and Nrh1 to investigate whether either of these sequences contain a proline residue that would allow them to unwind from an α-helix to a β-strand, as was observed for Gurken. Interestingly, while the TMD of zebrafish p75^NTR^ contains a proline close to the N-terminal end, zebrafish Nrh1 does not contain a proline within its TMD (Fig 6).

**Fig 6.**
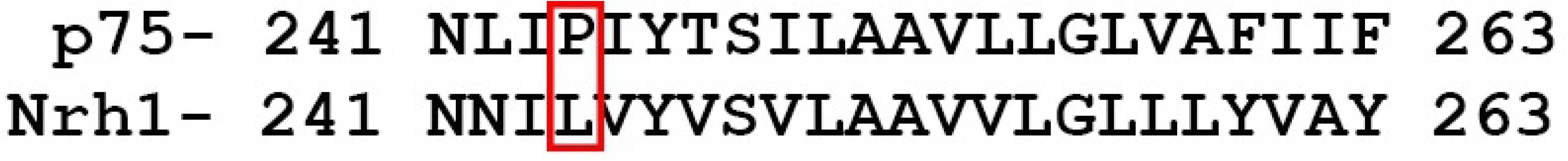
Alignment of the zebrafish p75^NTR^ and Nrh1 transmembrane domains. Red box indicates the proline residue in p75^NTR^ that is absent in Nrh1.

### Conclusions and Future directions

In this study we identified the *p75^NTR^* and *Nrh1* orthologues in zebrafish suitable for design of an assay system in which to test γ-secretase cleavage of these proteins *in vivo*. We observed that, while cleavage of zebrafish p75^NTR^ by γ-secretase is sensitive to DAPT, zebrafish Nrh1 is not sensitive to this γ-secretase inhibitor. This finding is consistent with a previous study in which human p75^NTR^ and Xenopus NRH1 were investigated *in vitro* [8]. Furthermore, our analysis of a chimeric Nrh1 protein in which the Nrh1 transmembrane domain is replaced by that of p75^NTR^ revealed that this domain alone is insufficient to confer γ-secretase cleavage susceptibility. This is not completely unexpected as there is evidence to suggest that other factors are crucial for selection/cleavage by the γ-secretase complex [5]. A greater understanding of the specificity of γ-secretase substrate selection might be reached by extending the chimaeric approach by exchanging each of the domains of these proteins. Indeed, a study (that we discovered while writing this manuscript) using a similar domain swap approach but with two unrelated type 1 transmembrane proteins (the non γ-secretase substrate Itgβ1 and γ-secretase substrate vasorin) found that both a permissive transmembrane and a permissive intracellular domain were required for γ-secretase cleavage, supporting our findings [31].

Other than the prerequisites of being a type 1 membrane protein and of shedding the ectodomain, it has previously been suggested that dimerisation of substrates and/or the structure of substrate α-helices may regulate γ-secretase activity [5]. This was not investigated in this study but is something that should be considered for future experiments using this assay system. Previous studies of the p75^NTR^ dimerisation domain AxxxGxxA found that, while this domain is not essential for dimerisation, altering its structure to LxxxLxxA via mutational analyses reduces γ-secretase cleavage [5, 32]. It was thought that this might be due to a stabilisation of the α-helix, inhibiting γ-secretase access by preventing unravelling of the transmembrane domain [5]. Zebrafish p75^NTF^ also contains the AxxxGxxA domain. However, in zebrafish Nrh1 the second alanine is replaced by leucine (AxxxGxxL) and we suggest this may have a similar effect to the LxxxLxxA mutant. It would be interesting to perform site directed mutagenesis on the L256 residue of the zebrafish Nrh1 dimerisation domain to assess whether this is a key feature preventing its cleavage by γ-secretase.

Another question that can potentially be addressed using this assay is the identity of the enzyme performing intramembranous cleavage of Nrh1. Since Kanning *et al* (2003) established that NRH1 does in fact produce cleavage products when treated with the PKC activator, PMA, a deeper investigation into what cleaves this protein may offer insights into alternative cleavage pathways for such substrates [8]. There are four main classes of proteases that perform intramembrane proteolysis. The first encompasses the aspartic proteases, including Presenilins (PSENs, the active subunit of γ-secretase), signal peptide peptidases (SPPs) and SPP-like proteases [33–35]. While γ-secretase cleaves proteins with type 1 membrane topology (C-termini towards the cytosol), the SPP and SPP-like proteases cleave proteins with type 2 membrane topology (N-termini towards the cytosol). Another class of I-CliPs consists of the Site-2 protease (S2P) and S2P-like proteases [36]. These are members of the metalloprotease family and cleave type 2 transmembrane proteases. Finally, there are the Rhomboid proteases, which are serine proteases [37]. Rhomboid proteases are highly specific in their substrate selection. The substrates selected are mostly type 1 transmembrane proteins, although there is also evidence to suggest they may cleave type 2 and multi pass membrane proteins in some cases [38, 39]. If we assume that NRH1 is a type 1 transmembrane protein like its homologue p75^NTR^, then we can reasonably exclude two of the above classes of membrane cleaving proteases as candidates, namely, SPP (and SPP-like) and S2P (including S2P-like). However, the orientation of NRH1 within the membrane has not yet been investigated. Our assay could be used to test a range of protease inhibitors to identify which enzyme(s) cleave Nrh1.

The question of the effect of the α-helix structure of p75^NTR^ or NRH1 TMD on their cleavage susceptibility has not yet been investigated. Previous observations from a study of the effects of the TMD of Gurken found that a β-sheet conformation was required for cleavage by the archaeal PRESENILIN homologue, MCMJR1. This raised the question of whether NRH1 can interact with PRESENILIN, but perhaps due to its TMD being in an α-helical conformation, cannot be cleaved by it. A simple amino acid sequence alignment of the zebrafish p75^NTR^ and Nrh1 TMDs (Fig 6) revealed that, while the p75^NTR^ TMD carries a proline residue that would supposedly allow it flexibility to conformationally change between an α-helix and β-sheet, the Nrh1 TMD lacks this residue. Interestingly, it has previously been observed that insertion of a single proline into a TMD can trigger cleavage of normally un-cleavable TMD’s [30]. It may, therefore, be of future interest either to insert or substitute a proline residue into the Nrh1 TMD to investigate the effect on its γ-secretase cleavage susceptibility. The results of such an experiment using our established assay may provide an answer for why Nrh1 is not naturally a γ-secretase substrate, while also contributing to the understanding of γ-secretase cleavage susceptibility. Regarding the previously observed cleavage of *Xenopus* NRH1 within its TMD and the question of what protease might be responsible for this cleavage, it has been observed that all intramembrane cleaving proteases (iCLIPs) prefer to cleave TMDs in their β-strand conformation [40]. If NRH1 is unable to enter this conformation, perhaps there is some other unknown enzyme responsible for this cleavage. If we wish to understand the cleavage properties of NRH1, it will be important to further investigate the conformational state of its TMD.

## Supporting information captions

**Figure S1. Duplication of the p75^NTR^ gene on chromosome 12 of *Danio rerio*.** Searching for p75^NTR^(ngfr) within the HomoloGene database on NCBI (https://www.ncbi.nlm.nih.gov/homologene/1877) returned the values displayed in this figure. The last two genes in the list are duplicate genes.

**Figure S2. Duplication of p75^NTR^ gene on chromosome 3 of Danio rerio, as of GRCz11.** The amino acid sequence of human p75^NTR^ was used in a tblastn search of the zebrafish genome (constrained to RefSeq_RNA). The two returned genes (indicated by the red box) represent the revised position of the p75^NTR^ duplicate gene on chromosome 3.

**Figure S3. Transmembrane domain predictions for Nrh1 and p75^NTR^ using TMHMM.** Plot of probabilities generated by TMHMM 2.0. A) p75^NTR^ TMD prediction B) Nrh1 TMD prediction. “outside” refers to the prediction that the sequence sits on the cytosolic side of the membrane and “inside” refers to the prediction that the sequence sits on the non-cytosolic (lumenal) side of the membrane. “transmembrane” refers to predicted transmembrane helices in the sequence.

**Table S1. Names, NCBI gene names and NCBI accession numbers of all genes used in phylogenetic studies**

**Table S2. Intensity ratios from western immunoblots for Figure 3**

**Table S3. Intensity ratios from western immunoblots for Figure 4**

**Table S4. Intensity ratios from western immunoblots for Figure 5**

**Text File S1. Construct overviews and full sequences**

